# A Kinesin-3 Recruitment Complex Facilitates Axonal Sorting of Enveloped Alpha Herpesvirus Capsids

**DOI:** 10.1101/704718

**Authors:** Julian Scherer, Ian B. Hogue, Zachary A. Yaffe, Nikhila S. Tanneti, Benjamin Y. Winer, Michael Vershinin, Lynn W. Enquist

**Author notes:** Correspondence should be addressed to (LWE).

## Abstract

Axonal sorting, the controlled passage of specific cargoes from the cell soma into the axon compartment, is critical for establishing and maintaining the polarity of mature neurons. To delineate axonal sorting events, we took advantage of two neuroinvasive alpha-herpesviruses. Human herpes simplex virus 1 (HSV-1) and pseudorabies virus of swine (PRV; suid herpesvirus 1) have evolved as robust cargo of axonal sorting and transport mechanisms. For efficient axonal sorting and subsequent egress from axons and presynaptic termini, progeny capsids depend on three viral membrane proteins (Us7 (gI), Us8 (gE), and Us9), which engage axon-directed kinesin motors. We present evidence that Us7-9 of the veterinary pathogen pseudorabies virus (PRV) form a tripartite complex to recruit Kif1a, a kinesin-3 motor. Based on multi-channel super-resolution and live TIRF microscopy, complex formation and motor recruitment occurs at the trans-Golgi network. Subsequently, progeny virus particles enter axons as enveloped capsids in a transport vesicle. Artificial recruitment of Kif1a using a drug-inducible heterodimerization system was sufficient to rescue axonal sorting and anterograde spread of PRV mutants devoid of Us7-9. Importantly, biophysical evidence suggests that Us9 is able to increase the velocity of Kif1a, a previously undescribed phenomenon. In addition to elucidating mechanisms governing axonal sorting, our results provide further insight into the composition of neuronal transport systems used by alpha-herpesviruses, which will be critical for both inhibiting the spread of infection and the safety of herpesvirus-based oncolytic therapies.

**Author Summary:** Alpha-herpesviruses represent a group of large, enveloped DNA viruses that are capable to establish a quiescent (also called latent) but reactivatable form of infection in the peripheral nervous system of their hosts. Following reactivation of latent genomes, virus progeny are formed in the soma of neuronal cells and depend on sorting into the axon for anterograde spread of infection to mucosal sites and potentially new host. We studied two alpha-herpesviruses (the veterinary pathogen pseudorabies virus (PRV) and human herpes simplex virus 1 (HSV-1)) and found viral membrane proteins Us7, Us8, and Us9 to form a complex, which is able to recruit kinsin-3 motors. Motor recruitment facilitates axonal sorting and subsequent transport to distal egress sites. Complex formation occurs at the trans-Golgi network and mediates efficiency of axonal sorting and motility characteristics of egressing capsids. We also used an artificial kinesin-3 recruitment system, which allows controlled induction of axonal sorting and transport for virus mutants lacking Us7, Us8, and Us9. Overall, these data contribute to our understanding of anterograde alpha-herpesvirus spread and kinesin-mediated sorting of vesicular axonal cargoes.

## Introduction

Neuronal cells establish and maintain polarity between the somatodendritic and axonal compartments via selective microtubule (MT)-based vesicle transport (1–3). Vesicles are propelled by opposing motor proteins of the cytoplasmic dynein and kinesin families towards either the MT minus ends or plus ends, respectively (4). The microtubules in axons are oriented predominantly with plus ends towards the axon terminus (5), and kinesin motors generally move cargoes in the anterograde direction, towards the plus end (6). Therefore, kinesin motors are thought to play a dominant role in sorting cargoes for axonal transport. Genetic screens have identified some of the kinesins that selectively transport cargoes across the axon initial segment (AIS) and into the axon (7). However, it is currently unknown what roles different kinesins, opposing dynein motors, MT modifications, MT-associated proteins, and the physical restrictions imposed by the actin-rich structure of the AIS play in axonal sorting processes (8–10). In this report, we studied the alpha-herpesviruses herpes simplex virus 1 (HSV-1) and pseudorabies virus (PRV; suid herpesvirus 1), robust cargos of MT-dependent vesicular axonal transport (11–13). PRV particle egress is a complex, multi-step process (14–16): newly-assembled progeny capsids are assembled inside the nucleus, cross the bilayers of the nuclear envelope, and gain access to the cytoplasm as unenveloped particles. In the cell bodies, these particles are transported towards the trans-Golgi network (TGN), where they complete assembly of the viral tegument protein layer, and acquire a double membrane by budding into an intracellular vesicle. The outer vesicle membrane is shed during the process of exocytosis at the plasma membrane, and the inner membrane serves as the virion envelope that mediates cell entry during the next infectious cycle. Importantly, egress vesicles must have the ability to interact with plus end-directed kinesin motors to move to the plasma membrane, in both neurons and non-neuronal cells. A subset of these vesicles must also recruit the kinesin-3 motor, Kif1a, to be sorted into the axon, cross the AIS, and move inside the axon towards distal egress sites (17, 18). We recently described that newly-assembled PRV particles in egress vesicles appear to move exclusively in the anterograde direction in axons, a behavior distinct from all other known axonal cargoes (13). Three PRV proteins encoded in a gene cluster (Us7 (glycoprotein I; gI), Us8 (glycoprotein E; gE), and Us9) are required for axonal sorting, and appear to function by recruiting Kif1a to the axonal egress vesicle (17, 19, 20). Us7 and Us8 encode for glycosylated multifunctional PRV membrane proteins with large cytoplasmic and ecto-domains (21), non-glycosylated Us9 is a small type II membrane protein that appears to function specifically in particle sorting and transport in neurons (22–25). These proteins are also conserved with HSV-1, where they play similar, but possibly non-identical, roles. Here, we use biochemical, biophysical, and microscopic techniques to determine how Us7-9 proteins form a tripartite Kif1a recruitment complex, and demonstrate that Kif1a recruitment to virion transport vesicles is sufficient for axonal sorting and transport. This study contributes to our understanding of the basic cell biology of axonal sorting and transport, host-virus interactions in the nervous system, as well as practical knowledge for the design of better therapeutic antiviral agents, vaccines, and herpesvirus-based gene therapy and oncolytic virotherapeutics.

## Results

### Anterograde spread efficiency of PRV US7-9 mutants and trans-complementation

We used a well-established *in vitro* assay of axonal sorting and subsequent anterograde spread of PRV from cultured neurons to epithelial cells (26). Primary rat superior cervical ganglia (SCG) neurons were cultured for two weeks in modified Campenot tri-chamber rings to separate cell bodies (soma compartment, S) from axon termini (neurite compartment, N) and virus-amplifying transformed epithelial indicator cells (Figure 1 A). To compare the influence of Us7, Us8, and Us9 on axonal sorting, transport and defects in kinesin motor recruitment, we infected the neuronal cell bodies with a panel of PRV mutant viruses and measured plaque-forming titer in the S and N compartments at 24h post-infection (hpi). Titers of S compartment lysates was a measure of efficient neuronal replication and were comparable to wild-type virus for all tested mutants (Figure 1 B-F). In parallel, titers of lysates of the epithelial cells in the N compartment represented the extent of anterograde axon-to-cell spread. For all PRV mutants, we also measured functional trans-complementation by transducing SCG neurons prior to PRV infection with non-replicating adenovirus vectors (AdV) encoding wild-type PRV proteins. The N compartment titer of PRV mutants lacking Us7 (ΔUs7) was 3.5 log units lower than the wild type titer (Figure 1 B). The ΔUs7-CT deletion mutant had a similar spread defect suggesting that the cytoplasmic C-terminal domain (CT) was required for efficient spread. The spread defects of mutants containing complete and partial Us7 deletions were all rescued by expressing wild-type Us7 via AdV transduction prior to infection (Figure 1B). PRV mutants lacking Us8 (ΔUs8) had a marked reduction of anterograde spread at 24hpi. While the CT of Us7 affects anterograde spread, deletion of the cytoplasmic C-terminal domain of Us8 had little effect on virus spread. By contrast, deletion of the N-terminal ecto-domain (NT) was as defective for anterograde spread as the complete deletion of Us8 (Figure 1 C). The spread defects of all Us8 mutants were rescued by AdV transduction of wild-type Us8. As shown previously, ΔUs9 PRV mutants were completely defective in anterograde spread at 24 hrs (Figure 1 D). Importantly, the spread defect of ΔUs9 was rescued by AdV transduction with wild-type Us9, but not by transduction of the YY(49, 50)AA di-tyrosine Us9 mutant (Us9-YY), consistent with earlier data (27, 28). We also tested other Us9 mutant proteins in AdV transduction studies for their capacity to complement the spread defect of the complete ΔUs9 mutant. A penta-serine Us9 mutant (Us9-5S) in which five identified phosphorylated serine residues (positions 38, 46, 51, 53, and 59) (17) were substituted with alanine failed to rescue the spread defect of ΔUs9 PRV. Similarly, a truncated Us9 protein that lacked the first 29 residues of the Us9 cytoplasmic N-terminal domain (Us9-N30) showed no complementation, whereas truncation of the first 21 residues (Us9-N21) showed weak complementation (∼2 log units).

**Figure 1.**
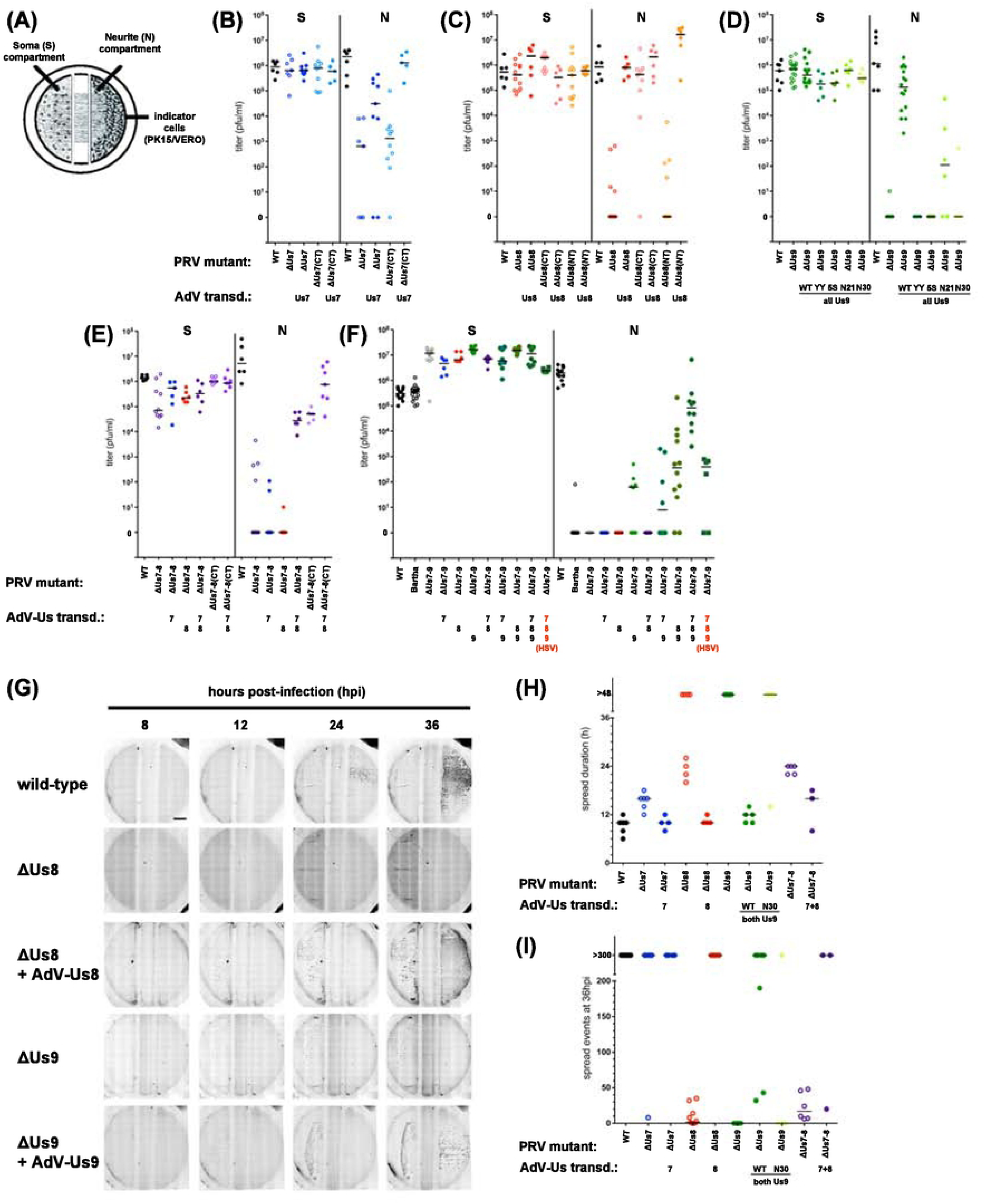
(A) Schematic representation of the modified Campenot trichamber culture system that physically separates SCG cell bodies (soma, S) from axons (neurite, N). For anterograde spread assays, SCG cells were infected with herpesviruses in S and indicator cells in N amplified spreading virus. Rescue of herpesvirus deletion mutants was via adenovirus transduction 2-3 days before infection. (B – D) Titers of PRV single Us gene deletion mutants 24 hpi and rescue by adenovirus transduction. Titers left and right of the solid vertical line represent S and N compartment titers, which indicate replication and anterograde spread, respectively. Each data point represents one chamber; horizontal bars indicate median values for each condition. (E) Titers of PRV ΔUs7/Us8 double mutant (ΔUs7-8) and rescue as described above. (F) Titers of PRV Bartha and ΔUs7/Us8/Us9 triple mutant (ΔUs7-9, PRV BaBe) and rescue as described above. Higher S chamber titers for PRV-ΔUs7-9 were due to a 5-fold increased pfu in the inoculum to facilitate replication for this virus. (G) Individual movie frames of representative anterograde spread chambers containing SCG neurons, which were infected with PRV strains as above. However, strains now also encode for mRFP-VP26 to indicate virus replication by red fluorescence, which was assessed every 2 h by large image microscopy starting at 6-8 hpi. Frames taken at 8, 12, 24, and 36 hpi are shown. All conditions showed red fluorescence in the S compartment 12 hpi, but only wild-type and rescue conditions of ΔUs8 and ΔUs9 viruses showed replication in N compartments. (H) Quantification of the spread duration determined by the elapsed time between detectable red fluorescence in the S and N compartments. Each data point represents one chamber; horizontal bars indicate median values for each condition. (I) Quantification of spread events in the N compartment at 36 hpi. Individual fluorescent foci were counted for the indicated conditions.

Anterograde spread of PRV-Us7/Us8 double deletion viruses (ΔUs7-8) was completely abrogated (Figure 1 E) and AdV trans-complementation was seen only when both proteins were expressed, but not with individual Us7 or Us8 expression. PRV mutants expressing Us7 and Us8 proteins both lacking their cytoplasmic C-termini (ΔUs7-8(CT)) showed an intermediate spread defect, indicating that deleting the cytoplasmic domains of both proteins had neither an additive nor synergistic effect.

Altogether, these data indicate that all three proteins have distinct roles in anterograde spread. For Us7, the relative minor contribution is mediated by the cytoplasmic domain, whereas for Us8, only the ecto-domain is required for efficient anterograde spread. Heterodimer formation of Us7 and Us8, which occurs via their ecto-domains, has been shown to be important for anterograde spread (19). To confirm the role of the Us7-9 proteins and further test the functionality of the transduced PRV proteins, we used AdV transduction to complement the severe anterograde spread defect of a PRV mutant lacking all three genes (PRV BaBe; ΔUs7-9) (Figure 1 F). PRV BaBe has spread defects comparable to the parential PRV vaccine strain ‘Bartha’ *in vivo* and *in vitro* (22, 28). AdV expressing Us7, Us8, or both in combination failed to complement the spread defect of PRV BaBe. However, expressing Us9 alone or in combination with either Us7 or Us8 showed modest complementation. However, expression of all three proteins with AdV restored efficient anterograde spread to PRV BaBe as efficiently as complementing single-mutants (Figure 1 B-D).

Finally, we determined if the homologous Us7-9 proteins of herpes simplex virus 1 (HSV-1) could complement the spread defects of PRV Us7-9 mutants. AdV transduction of all three HSV-1 Us7-9 proteins together were modestly effective in restoring PRV BaBe anterograde spread; similar to the extent of complementation of the PRV BaBe defect by PRV Us8/Us9. Complementation of the PRV BaBe spread defect by single HSV-1 Us8 or Us9 proteins was not observed (Suppl Figure 1 A). However, we found that the HSV-1 Us7 protein was able to complement the spread defect of the PRV-ΔUs7 virus, but to a much lower extent than the PRV Us7 protein. We note that the HSV-1 wild-type virus had a marked slower anterograde spread phenotype compared to PRV (Suppl Figure 1 B). Interestingly, only HSV-1 Us7, but not HSV-1 Us8 or Us9, show complementation of the corresponding PRV single mutant virus, however with slow kinetics resembling HSV-1, but not PRV, wild-type.

### The kinetics of limited anterograde spread promoted by deletions of Us7, Us8, or Us9

The amount of time taken to spread from axons to epithelial cells is a measure of the extent of axonal sorting and axonal transport of virions in neurons, as well as the timing and extent of axonal egress to the amplifying epithelial cells. To determine the kinetics of limited anterograde spread observed after neuronal infection by PRV mutants with deletions of Us7, Us8, or Us9, we constructed mutants also expressing the mRFP-VP26 capsid fusion protein (29). This fusion protein enabled us to visualize infection of neurons and indicator cells using whole chamber fluorescence microscopy as well as facilitating tracking of individual particles. We routinely performed time-lapse imaging of infected SCG neuronal cultures continuously from 8 – 48 hpi, in some cases up to 72 hpi (Figure 1G-H and data not shown). PRV wild-type and all tested mutants produced significant red fluorescence in the S compartment at ∼10 hpi. Differences in the kinetics of anterograde spread among mutant viruses became apparent when we measured the elapsed time between the appearance of red fluorescence in the S compartment and subsequent detection in epithelial cells in the N compartment (Figure 1 H). On average, wild-type PRV took 10 h to spread from the S compartment to the indicator cells in the N compartment. The ΔUs7 mutant took 16 h. For the ΔUs8 mutant, several replicates had detectable spread that was delayed by 10-16 h, and several had no spread visible after 48 h or longer. This finding of a mixed spread phenotype for Us8 mutants is reminiscent of the mixed spread defects for Us8 mutants observed *in vivo* (30). Interestingly, the spread defect of the ΔUs7/ΔUs8 double mutant was intermediate, between the defects seen for either single mutants, again indicating there is no additive or synergistic effect of deleting both proteins. ΔUs9 virus mutants had no detectable spread to epithelial cells (Figure 1H), even up to 72 hpi. All spread defects could be complemented by AdV transduction of wild-type PRV proteins. We also counted the number of fluorescent capsid foci in the N compartment epithelial cells at 36 hpi (Figure 1G). PRV wild-type and all AdV-complemented conditions resulted in similar foci counts (>300 foci), as did infection with the ΔUs7 mutant. However, infection with the ΔUs8, ΔUs7/ΔUs8 and ΔUs9 mutants showed a markedly reduced number of foci in the N compartment epithelial cells (0–50). These data indicated that Us7-9 mutations have different effects on the kinetics of anterograde spread. When compared to the spread of wild type PRV, ΔUs9 mutants showed no anterograde spread at all, spread of the ΔUs8 mutant was markedly reduced, and spread of the ΔUs7 mutant showed an intermediate delay.

### Analysis of post-translational Us7-9 modifications on anterograde spread

The presence of Us9 protein is critical for anterograde spread, and phosphorylation of Us9 appears to be critical as well. The penta-serine mutation that removes Us9 phosphorylation sites is unable to restore wild type spread kinetics to the ΔUs9 mutant. It has also been established that maturation of the glycoproteins Us7 and Us8 in the Golgi compartment is required for their function (21, 31, 32). We next analyzed post-translational events by Western blotting of infected cell lysates with mutant strains (Figure 2A). We cultured SCG neurons for two weeks, infected the cultures at an MOI of 10 with PRV wild-type or mutant strains for 6, 12, or 24 h, and analyzed the cell lysates by Western blot. Treatment with the drug Brefeldin A (BrefA), which inhibits ER-to-Golgi trafficking and protein glycolysation at the Golgi (33), served as a control. As previously reported, Us7 maturation did not occur in ΔUs8-infected cells, whereas Us8 maturation was only partially delayed in ΔUs7-infected cells (31). Of note, cells infected with the ΔUs7(CT) mutant, which lacks the cytoplasmic tail of Us7, exhibited normal Us8 maturation, and cells infected with the ΔUs8(CT) mutant, which lacks the cytoplasmic tail of Us8, exhibited normal Us7 maturation, indicating that the cytoplasmic tails are not required for proper glycoprotein maturation. Importantly, Us9 expression and phosphorylation occurred in all Us7 or Us8 mutant infected or BrefA-treated cells similar to wild-type PRV infections. Since BrefA has strong effects on Us7 and Us8 maturation, we also investigated the kinetics of BrefA-induced Golgi disruption on efficiency of anterograde spread (Figure 2B). BrefA was applied only to the S compartment neuronal cell bodies at the indicated times post infection. Early addition of the drug at 2.5 hpi inhibited viral replication as indicated by strongly reduced S compartment titers (32). Later addition of the drug at 5 hpi had minor effects on replication but essentially eliminated spread to indicator cells in the N compartment. Addition of BrefA to the S compartment at 8.5 hpi resulted in minimal replication and spread defects, suggesting that the first progeny particles are sorted into the S compartment and spread to detector cells between 5-8.5 hpi, consistent with our live imaging results (Figure 1 H). These results indicate that BrefA treatment affects axonal sorting and anterograde spread late in infection. In sum, post-translational modifications of all three PRV proteins play important roles in anterograde spread.

**Figure 2.**
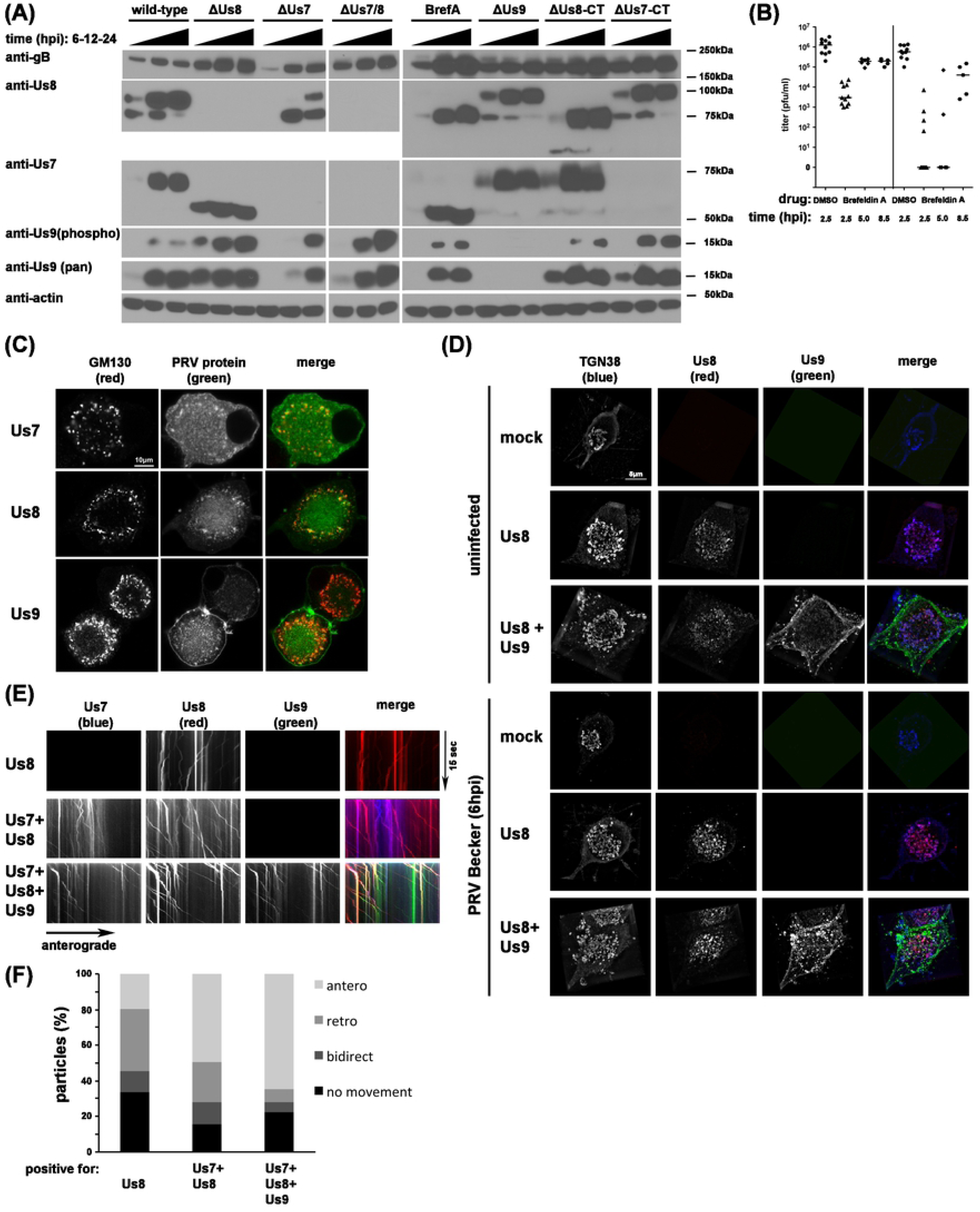
(A) Western blot of SCG cell lysates illustrating the maturation process of PRV glycoproteins. SCG neurons were infected with PRV strains, lysates were taken at 6, 12, and 24 hpi, and processed for Western blotting using anti-Us7, anti-Us8, and phosphor- and pan-anti-Us9 antibodies. Antibodies against gB or actin were used as infection or loading control. Us7 and 8 maturation was visible by band shift in the wild-type infected neurons; the lower band represents the ER species and the higher band the mature, glycosylated Golgi form of these proteins. Us7 maturation was abolished for ΔUs8 viruses and Us8 maturation was severely inhibited in ΔUs7 viruses. Deletion of Us9 had no effect on Us7 or Us8 maturation. Conversely, deletion of Us7 and Us8 or disruption of Golgi function with Brefeldin A (BrefA) had no effect on Us9 expression and phosphorylation. Importantly, the Us8 N-terminus confers the ΔUs8-mediated defect in Us7 maturation since the Us8 C-terminal deletion ΔUs8(CT) shows mature Us7. Similarly, Us8 matures with wild-type kinestics in ΔUs7(CT) infected cells. The Us7 antibody recognizes the C-terminus of the protein and cannot detect ΔUs7(CT). (B) Titers of PRV wild-type infected anterograde spread assays at 24 hpi as described in Figure 1, but in Brefeldin A (BrefA) treated cells. SCG neurons were infected with PRV wild-type and treated with Brefeldin A at either 2.5, 5, or 8.5 hpi, DMSO treatment at 2.5 hpi as control. (C) Confocal imaging of SCG neuronal cell soma after transduction with indicated PRV proteins. Cells were permeabilized and stained with the Golgi marker GM130. (D) Super-resolution imaging of SCG soma using structured illumination microscopy (SIM). Cells were transduced with the indicated PRV proteins and fixed either uninfected or after 6 h of PRV Becker infection. Cells were permeabilized and stained with the trans-Golgi marker TGN38. Representative images are shown in 3D reconstruction mode. (E) TIRF microscopy of live SCG axons grown in compartmentalized cultures. Cells were transduced with indicated PRV proteins and imaged at ∼12frames/s in three-color mode. Axonal co-transport of Us7-9 was observed without PRV infection. (F) Quantification of the directionality of the moving particles observed in (E).

### Complex formation and motor recruitment occurs at the trans-Golgi network

We next determined if the AdV-transduced proteins maintain the capacity to localize to the Golgi on their own, in the absence of PRV infection (Figure 2 C). Confocal imaging of SCG neurons transduced with Us7-mTurquoise2, Us8-mCherry, or GFP-Us9 all show colocalization of the PRV proteins with the Golgi marker GM130. Us8-mCherry most strongly colocalized with GM130, whereas GFP-Us9 also exhibited strong vesicular and plasma membrane localization, and the Us7-mTurquoise2 fusion protein also showed a diffuse signal throughout the soma. We used super-resolution microscopy to determine if Us8 and Us9 are in close proximity at the Golgi compartment (Figure 2 D). We observed weak colocalization between Us8-mCherry and GFP-Us9 at the trans-Golgi compartment (as marked by TGN38) in uninfected cells. However, the colocalization was greatly enhanced after infection with wild type PRV Becker (17). Interestingly, Us8 showed significant colocalization with TGN38 under both conditions. These results indicate that in the course of a wild-type infection, Us7-9 localize to the Golgi, where they are present in close proximity. In fixed cell analysis, we also observed discrete colocalized puncta of AdV-transduced Us7-9 along the axons of uninfected SCG neurons (data not shown). Subsequent live cell TIRF microscopy of SCG cell cultures grown in Campenot chambers (13), transduced with either Us8-mCherry alone or in combination with Us7-mTurquoise2 and GFP-Us9, revealed the dynamic behavior of these puncta (Figure 2 E and F). In the axons of uninfected neurons, Us8 alone localized to individual motile vesicles that exhibited bidirectional retrograde and anterograde movements. When Us7 and Us8 were co-expressed, they colocalized in motile vesicles, but exhibited a greater proportion of anterograde movement. When all three proteins, Us7-9 were co-expressed, distinct puncta positive for all three proteins showed the highest percentage of anterograde processive motility. Similar experiments with HSV-1 Us7-9 also revealed that all three viral proteins moved together in the absence of viral infection (Suppl Fig 1 C). These data indicate that the specific anterograde-only motility pattern observed for PRV egress vesicles (13) is very closely mimicked by the three Us7-9 proteins, even in the absence of infection. They further support results that Us7-9 of HSV-1 and PRV have the capacity to induce anterograde vesicle motility.

### PRV infection promotes robust Us7-9 complex formation

Given that Us7-9 colocalize at the trans-Golgi network and move together in axons, we attempted to characterize the three protein complex after infection (Figure 3 A). We transduced PK15 cells with AdV encoding Us8-mTurquoise-HA, together with GFP-Us9, or with GFP-Us9 and Us7-mTurquoise-Flag. Cells remained uninfected (mock), or were infected with PRV Becker for 12 hpi, lysates were prepared, and immunoprecipitated with an anti-HA tag antibody. As expected based on previous reports, Us7 and Us8 form a strong interaction independent of PRV infection. However, we were unable to show an association with Us9 in transduced, but uninfected cells. However, Us9 was clearly present in immunoprecipitates of Us7-9 in PRV-infected samples. Untagged endogenous Us9 and GFP-tagged AdV-transduced Us9 both co-immunoprecipitated with the Us7/Us8 heterodimer. We note that in multiple repeats of this experiment, we were unable to detect cytoplasmic dynein or kinesin motor proteins by co-immunoprecipitation of Us7-9 (data not shown). We next turned to step gradients and co-sedimentation experiments as a gentler approach to identify possible interactions of the Us7-9 complex with motor proteins (Figure 3 B). Isolating detergent resistant membranes (DRM) from neuronal cell cultures by gradient ultra-centrifugation enriches for “lipid raft”-like membrane sub-compartments, which have been implicated in Us7-9 function and PRV axonal spread (34). We compared the presence of Kif5 (kinesin-1) and Kif1a (kinesin-3) in DRM fractions of mock, PRV wild-type, or ΔUs9-infected SCG neurons. Kif1a and Kif5 motors drive anterograde transport of cellular axonal cargoes, and have both have been reported to be involved in PRV and HSV-1 anterograde transport (17, 35, 36). In wild-type infected samples, we detected an increase in Kif1a in the DRM fractions compared to mock and ΔUs9-infected conditions, but no enrichment of Kif5. Cytoplasmic dynein was present at low levels in DRM fractions, but was independent of PRV infection or the presence of Us9 (data not shown). Taken together, this biochemical evidence indicates, that during PRV infection Us7-9 form a tripartite complex that specifically recruits Kif1a in a Us9-dependent manner, but this complex is easily disrupted by standard cell lysis methods.

**Figure 3.**
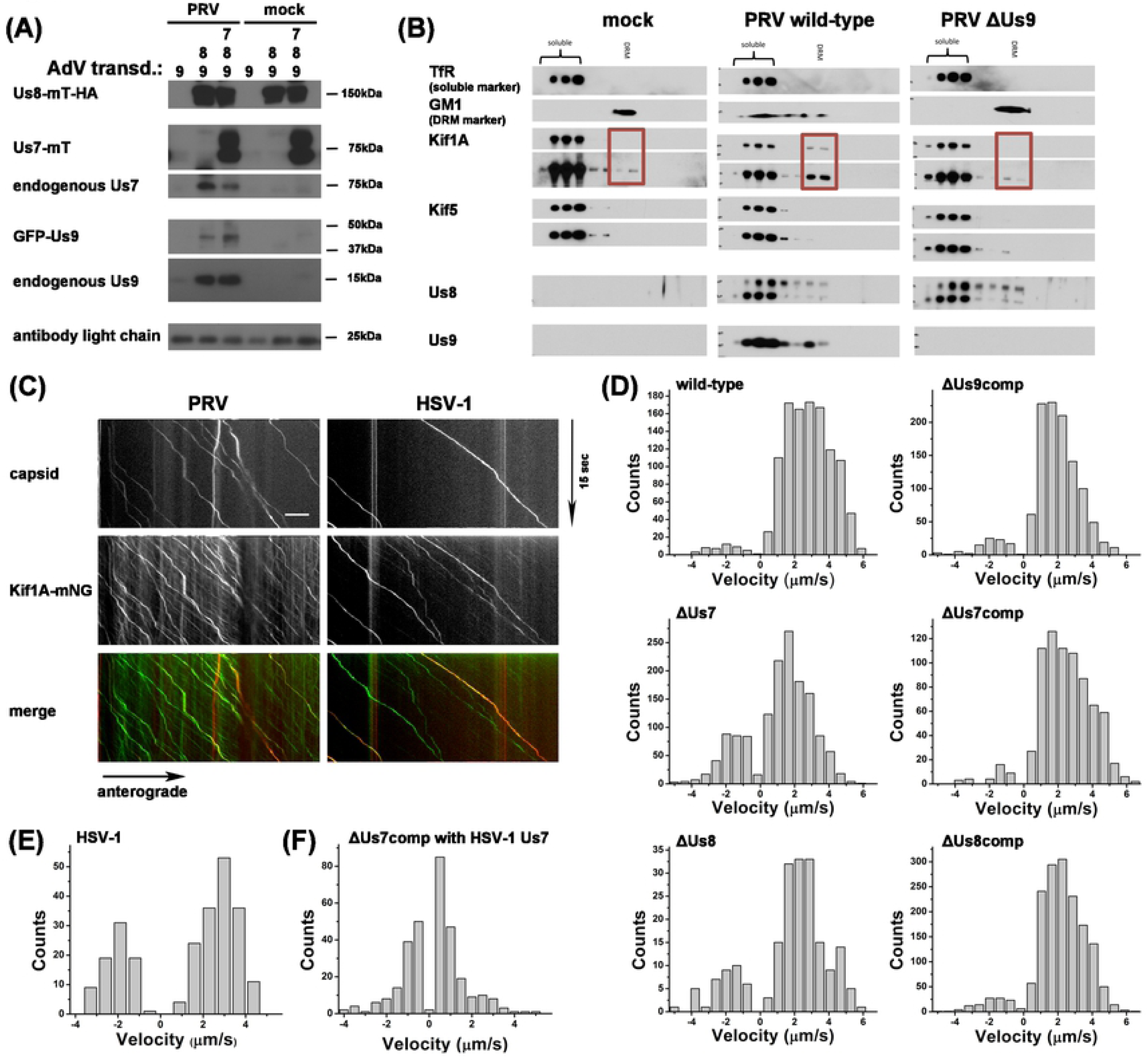
(A) Immunoprecipitates of anti-HA bead fractions of PK15 lysates transduced with mock (lanes 1 & 4), Us8-mT-HA and GFP-Us9 (lanes 2 & 5), or Us8-mT-HA, GFP-Us9, and Us7-mT (lanes 3 & 6) and either PRV infected (lanes 1-3) or uninfected (lanes 4-6). Imunoprecipitates were processed for Western blotting and the membrane was incubated with antibodies against HA-tag, GFP-tag, Us7, and Us9. (B) DRM fraction isolation from SCG neuronal cultures as described in [Lyman et al., PLoS Path, 2008] using mock infected cells or cells infected with wild-type or ΔUs9 PRV. Fractions were processed for Western blotting and membranes were incubated with antibodies against Kif1a, Kif5, Us8, Us9 and TfR (soluble marker) or Cholera Toxin B (CTX-B, DRM marker). (C) TIRF microscopy of live SCG axons grown in compartmentalized cultures. Cells were transduced with full-length Kif1a-mNG, infected with red capsid tagged PRV or HSV-1 and imaged at ∼19frames/s at 12-14hpi. Axonal co-transport of Kif1a was observed with both viral capsids. (D) Motility analysis of segment velocities during capsid egress as described in Scherer et al., JVI, 2016 for PRV wild-type, mutant, and compensation (comp) conditions. (E) Same as in (D) but for HSV-OK14 egress. (F) Same as in (D), but for ΔUs7 PRV compensation with HSV-1 Us7.

### Kif1a transports PRV and HSV-1 in axons with Us7 playing a critical role in velocity differences

To determine whether Kif1a motor proteins colocalize and co-transport with PRV and HSV-1 particles in axons, we again used live cell TIRF microscopy of SCG neurons grown in Campenot chambers (Figure 3 C). SCG neurons were transduced with AdV expressing full-length Kif1a-mNeonGreen (Kif1a-mNG), and co-infected with PRV or HSV-1 recombinants expressing mRFP-VP26 capsid tags (PRV 180 and HSV-OK14). We detected distinct co-translocation of Kif1a with fluorescent capsids expressed by both viruses. In similar experiments, we were unable to detect co-translocation with a fluorescent protein-tagged Kif5 (data not shown). To measure virus particle transport, we performed automated single particle tracking to extract particle velocities (Figure 3 D-F). For wild-type PRV, we detected almost exclusively anterograde motility, with a broad distribution of anterograde velocities up to 6 μm/sec (13). Since we were unable to detect any ΔUs9 particles in axons, we analyzed AdV-complemented ΔUs9 (ΔUs9comp) particle motility only. With AdV transduction to express GFP-Us9, we also observed almost exclusively anterograde motility, albeit with a reduced maximum velocity. One possible explanation for this reduction may be steric hindrance by the GFP tag on the transduced GFP-Us9, or that the quantity and kinetics of AdV transduced Us9 expression does not complement as efficiently as endogenous Us9 expression. PRV ΔUs7 particles had somewhat reduced maximum velocities, but also the appearance of more retrograde motility. These effects were rescued by AdV transduction of Us7 (ΔUs7comp). This finding suggests that Us7 might play a role in inhibiting cytoplasmic dynein-mediated retrograde transport and accelerating Kif1a motility during anterograde transport. The data also provides a possible explanation for the intermediate spread defect observed for ΔUs7 strains (Figure 1 B). In addition, we analyzed a small number of rare motile ΔUs8 particles in the axon. Similar to the ΔUs7 condition, ΔUs8 particles had fewer instances of high velocity anterograde transport, and the appearance of more retrograde motility, which was rescued by AdV transduction of Us8 (ΔUs8comp).

We extended our analyses to HSV-1 capsids in axons (Figure 3 E). We observed a single peak of velocity of HSV-1 particles in the anterograde direction, and also a distinct retrograde peak, implying a more extensive contribution of cytoplasmic dynein during axonal transport of HSV-1. These findings provide an explanation of why HSV-1 anterograde spread appears delayed compared to PRV spread in our system. Finally, we investigated the specific contribution HSV-1 Us7 to the slower motility profile of HSV-1 by complementing PRV-ΔUs7 with AdV expressing HSV-1 Us7 (Figure 3 F). We detected motile capsids in axons suggesting that HSV-1 Us7 can complement PRV-ΔUs7 (Suppl Figure 1 B). However, these complemented particles showed more retrograde motility and an apparent lack of high anterograde velocities, when compared to the PRV Us7 complementation motility profile. Taken together, these data strongly point to a role of PRV Us7 in reducing retrograde, cytoplasmic dynein-mediated transport events and initiating pathways that depend on Us9 to accelerate Kif1a-mediated anterograde transport.

### Artificial Kif1a recruitment is sufficient to rescue PRV mutants devoid of Us7-9

This report, together with previous publications, has shown that Us9 is necessary for recruitment of Kif1a, axonal sorting of PRV particles, and anterograde spread of PRV. To determine whether Kif1a recruitment is sufficient for anterograde sorting, transport, and spread, we constructed a system for drug-inducible Kif1a recruitment to intracellular virus particles, independent of Us9 (Figure 4A). We modified a previously-described drug-inducible motor recruitment system where two protein domains, FKBP and FRB, heterodimerize only in the presence of a membrane-permeable rapamycin analogue (37). We fused two FKBP heterodimerization domains to the cytoplasmic C-terminus of glycoprotein M (gM-FKBP), a viral membrane protein that is present in the virion envelope and surrounding intracellular transport vesicle (38), in PRV ΔUs9 or ΔUs7-9 genetic backgrounds. A construct consisting of Kif1a motor domain tagged with GFP and an FRB heterodimerization domain was expressed in neurons via AdV transduction. Upon addition of the heterodimerizer drug, FKBP and FRB interact, recruiting the FRB-tagged Kif1a motor to the virion egress vesicle containing gM-FKBP (Figure 4A). We found that SCG neurons supported anterograde spread of ΔUs9 mutant PRV only after Kif1a-GFP-FRB transduction and addition of the heterodimerizer drug (Figure 4 B). We also observed anterograde spread with a ΔUs7-9 mutant expressing gM-FKBP. These data indicate that artificial Kif1a recruitment to egress vesicles is sufficient for axonal sorting, transport, and anterograde spread, even in the absence of Us7-9. Next, we analyzed the induced motility of ΔUs9 and ΔUs7-9 particles by TIRF microscopy in the presence or absence of the heterodimerizer drug (Figure 4 C). Both ΔUs9 and ΔUs7-9 mutants showed anterograde capsid transport after addition of the heterodimerizer drug. Subsequent motility analysis (Figure 4 D) indicates slower, but anterograde-only velocities of ΔUs9 particles, whereas ΔUs7-9 particles also exhibited some movement in the retrograde direction. The retrograde motility of ΔUs7-9 particles is consistent with the phenotype of ΔUs7 and ΔUs8 particles (Figure 3 D). Overall, these results support a model suggesting that PRV-Us9 represents a<u>n</u> essential component in the tripartite Kif1a recruitment complex of PRV, which is able to modulate Kif1a speed to higher velocities, and/or reduce the function of slower anterograde and opposing retrograde motors. PRV Us7 and Us8 might also support Us9 in accelerating wild-type Kif1a motors (though not the artificial Kif1a-GFP-FRB motor) and inhibiting retrograde movement mediated by cytoplasmic dynein.

**Figure 4.**
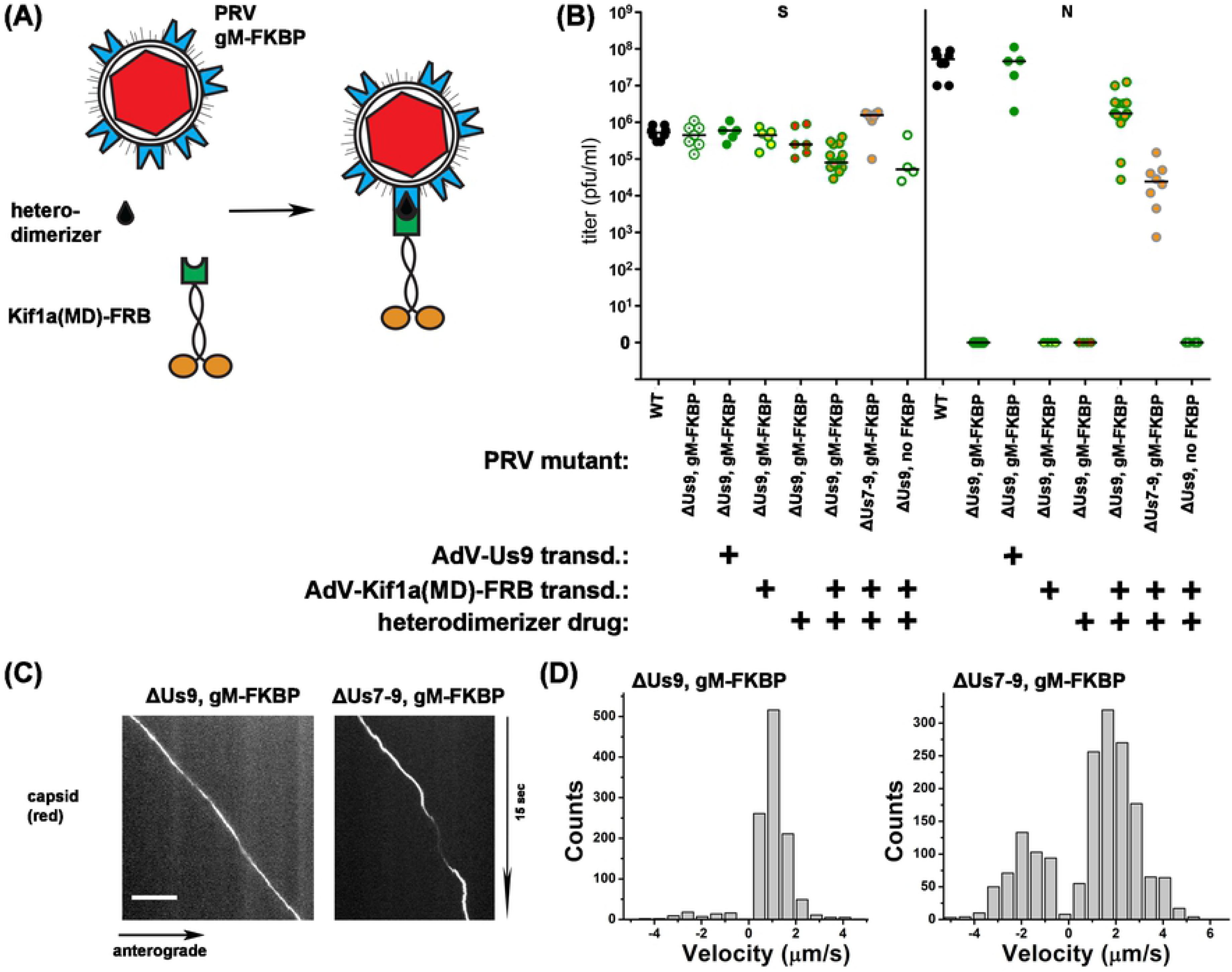
(A) Cartoon representation of the elements of the engineered inducible system for Us9-independent PRV spread. (B) Titers of indicated PRV mutants 24 hpi and rescue by adenovirus transduction with GFP-Us9 or Kif1a-FRB and addition of heterodimerizer drug. Titers left and right of the solid vertical line represent S and N compartment titers, which indicate replication and anterograde spread, respectively. Each data point represents one chamber; horizontal bars indicate median values for each condition. (C) TIRF microscopy of live SCG axons grown in compartmentalized cultures. Cells were transduced with Kif1a-FRB, infected with red capsid tagged gM-FKBP ΔUs9 PRV or ΔUs7-9 PRV and imaged at ∼19frames/s at 12-14hpi. (D) Motility analysis of segment velocities of (C).

### The Kif1a recruitment complex remains associated with egressing particles

Next, we determined if the tripartite Kif1a recruitment complex remains associated with the capsid during transport along the axon (Figure 5 A). We transduced SCG neuronal cultures in Campenot chambers with AdV expressing Us7-mTurquoise2 (Us7-mT), Us8-mTurquoise2 (Us8-mT), or GFP-Us9 and infected with the corresponding PRV deletion mutant expressing mRFP-VP26 to label capsids. At 12 hpi, we observed motile progeny capsids using live cell TIRF microscopy. In all conditions, the majority of observed motile particles showed distinct co-location of capsid and envelope proteins along the length of the run. The signal for Us7 and Us8 in the axon was almost exclusively associated with moving capsids, but puncta positive for Us9 were also observed to move independently of capsids. We also observed co-translocation of HSV-1 mRFP-VP26 capsids and AdV-transduced HSV Us7-mTurquoise2 and HSV GFP-Us9 (Figure 5 B). These data indicate that the triple complex of Us7-9 is closely associated with PRV and HSV-1 capsids undergoing anterograde transport in the axon of neuronal cells.

**Figure 5.**
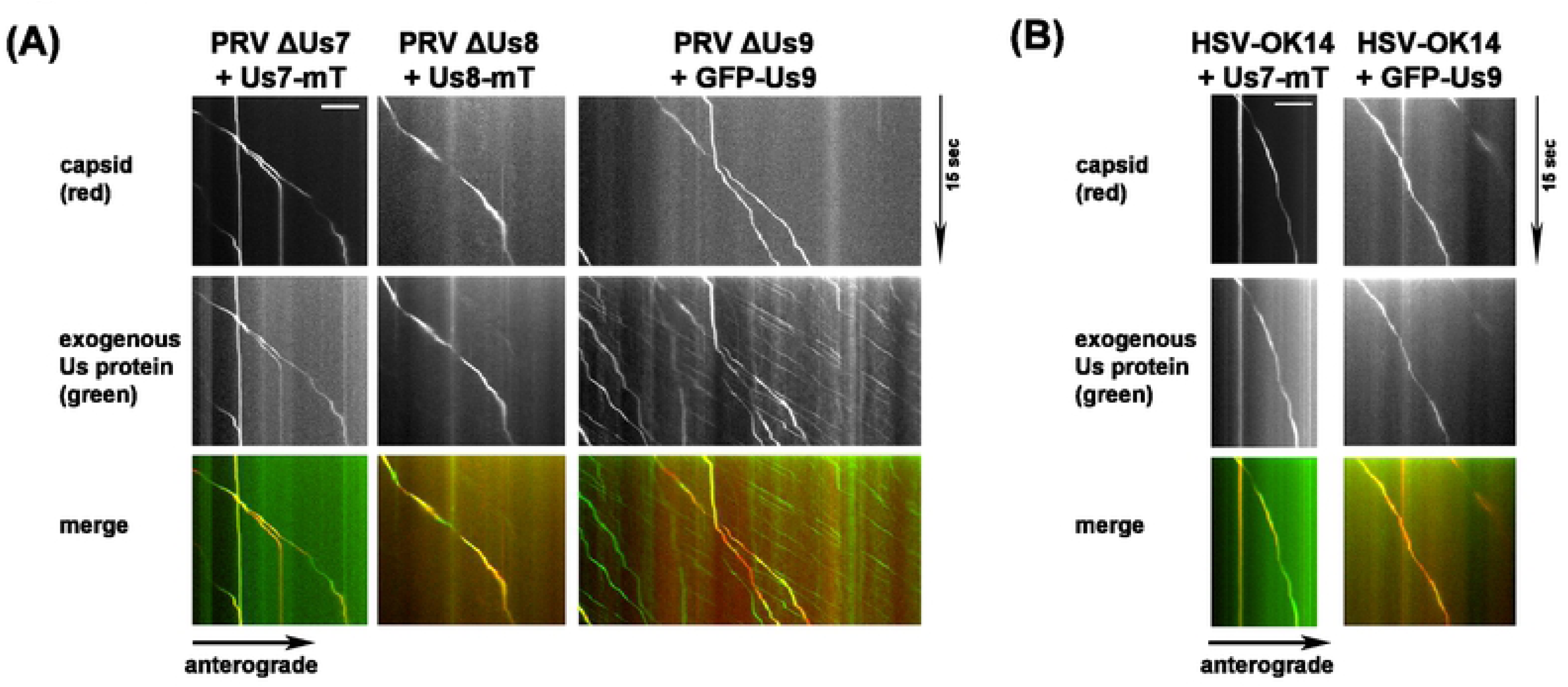
(A) TIRF microscopy of live SCG axons grown in compartmentalized cultures. Cells were transduced with indicated PRV proteins, infected with red capsid tagged PRV mutant lacking the corresponding protein and imaged at ∼19frames/s at 12-14hpi. Axonal co-transport of the viral envelope protein was observed with all viral capsids. (B) Same as in (A), but with transduction of HSV-1 proteins and red capsid tagged HSV-1 wild-type virus.

## Discussion

Here we show how a tripartite complex consisting of the PRV proteins Us7, Us8, and Us9 recruits the kinesin-3 motor protein Kif1a to viral egress vesicles. Kif1a recruitment alone - via Us7-9 or via an engineered, drug-inducible system - is sufficient to sort the virion egress vesicle into the axon, leading to anterograde transport, and spread to epithelial cells. Furthermore, super-resolution and TIRF microscopy, coupled with motility analysis at high temporal and spatial resolution, indicates that each member of the tripartite complex has distinct functions. We present evidence that Us8 has the strongest capacity to localize to the trans-Golgi network where it presumably recruits Us7 via tight heterodimer formation, leading to subsequent Us7 maturation. Us9 can localize to the Golgi independently of Us7/Us8 or full PRV infection. In order to function in axonal sorting and transport, the Us9 protein must be present at the trans-Golgi network, likely phosphorylated at critical serine residues, and form a tripartite complex with Us7 and Us8, potentially via the C-terminal domain of Us7. Since egress from cell bodies (not axons) occurs efficiently in the absence of Us9, and only a minority of egress vesicles sort into the axon late in infection, the kinetics of Kif1a-recruitment are not yet clear. The specific complex of Us9 with Us7 and Us8 may form only in a subset of secondary envelopment sites or egress vesicles (39), which are then destined to recruit Kif1a and be sorted to the axon. Based on the existence of fast moving vesicular structures positive for Us7-9 in axons of uninfected SCG neurons, Kif1a recruitment does not appear to require any additional viral factors aside from the Us7-9 complex itself.

All tested viral envelope proteins, as well as the Kif1a motor protein, remain tightly associated with the capsid during anterograde axonal transport, and can be observed along the entire length of the axon. Given the long run length of individual particles, we conclude that the dissociation rate of the Us7-9 recruitment complex and the motor protein is low. This conclusion is further supported through our biochemical evidence of a Us9-dependend Kif1a recruitment to “lipid raft” like membrane structures and through the motility analysis of engineered heterodimer-induced capsid transport. The FKBP-FRB interaction has a half-life of ∼18 h, and the resulting anterograde-only motility resembles that of wild-type particles. It remains unclear why the heterodimer-induced capsid structures do not have higher velocity runs (Figure 4D). Possibly, the lack of Us9 keeps the Kif1a motor operating at a lower velocity. Alternatively, the truncated Kif1a-GFP-FRB construct used for these experiments may move with a slower maximum velocity in general. Additionally, AdV transduction might not confer complete complementation, as may also be the case with ΔUs9 complementation by AdV-transduced GFP-Us9 (Figure 3 D). The role of Us8 appears to be in the initiation of stable complex formation at the trans-Golgi network via heterodimerization with Us7. Anterograde spread does not depend on the Us8 cytoplasmic domain, which rules it out as a likely contact interface with Us9 or Kif1a. However, potential interactions via the trans-membrane domains of the proteins cannot be ruled out. It is clear that the Us8 ecto-domain is critical for spread *in vitro* and *in vivo* (19, 40), which is presumably due to Us7 binding and also the ability to interact with tegument proteins during secondary envelopment (31). In this study, we found two potential roles for Us7. One is that Us7 may inhibit cytoplasmic dynein on transport vesicles and the other is that Us7 in complex with Us8 and US9 may modulate Kif1a acceleration. Such activities for Us7 may explain why wild-type HSV-1 progeny particles exhibit some retrograde motility in axons. HSV-1 Us7 lacks the capacity to inhibit cytoplasmic dynein on transport vesicles, which enables some retrograde transport of HSV vesicles. As a result, HSV-1 has slower anterograde spread kinetics than PRV, moves slower anterogradely inside the axon during egress, and that ΔUs7 PRV complemented with HSV-1 Us7 leads to slow moving egress capsids. The kinetics of spread of these complemented particles resemble HSV-1 wild-type particles. Therefore, it appears that Us7 in both alpha-herpesviruses, PRV and HSV-1, represents a critical factor that determines the kinetics of anterograde spread. The difference in Us7-mediated egress kinestics may indicate a host-specific divergence as alpha-herpesviruses have evolved to infect hosts of varying sizes and at different anatomical sites, which may require faster anterograde transport with less of a retrograde component. It will be interesting to compare PRV and HSV-1 axonal transport to that of other alpha-herpesviruses with different natural histories.

Our results indicate that both PRV and HSV-1 use a robust Kif1a recruitment mechanism for efficient axonal sorting and transport. We directly tagged the full-length Kif1a motor protein and showed co-transport in axons of individual PRV and HSV-1 proteins on egressing PRV and HSV-1 particles. (Figure 3C). Other studies have implicated other kinesin motors using more indirect methods, including truncated motor proteins or motor knock-down experiments (36). For PRV, motor recruitment alone appears to be necessary and sufficient to guide progeny capsids in enveloped structures into the axon process and subsequently along microtubules past the axon initial segment to distal egress sites. The interaction of the tripartite US7-9 recruitment complex with the Kif1a motor protein is required only for axonal sorting and subsequent axonal transport. This interaction is not essential for transport and egress of virus particles from non-neuronal cells. We speculate that interactions between the Us9 cytoplasmic tail, Us7, and Kif1a represent potential targets to block viral transport and spread in axons, even after viral replication has been initiated. In addition, these findings have implications for understanding of axonal sorting of cellular cargoes in uninfected cells, and underscore the importance of motor recruitment in axonal sorting.

## Materials & Methods

### Cell Lines

Cell lines used for this study, including human 293FT cells (Invitrogen), rat2 cells (ATCC #CRL-1764), African green monkey kidney epithelial cells (VERO, ATCC #CCL-81), and porcine kidney epithelial cells (PK15, ATCC# CCL-33), were maintained in DMEM (HyClone) supplemented with 10% FBS (HyClone) at 37°C and 5% CO2.

### PRV and HSV-1 Recombinant Viruses

Alpha-herpesvirus strains used in this study are summarized in Table 1. PRV Becker and HSV-1 17+ were used as wild-type strains. All virus titers were determined by serial dilution plaque assays on PK15 cells for PRV strains or VERO cells for HSV strains, and predicted Us7-9 gene expression was confirmed by Western blotting of virus infected cell lysates. PRV strains PRV740, PRV741, and PRV742 were constructed by co-infection of PRV180, expressing an mRFP-VP26 capsid tag, with PRV98 (Us7 null), PRV99 (Us7/Us8 null), and PRV107 (Us8-CT deletion), respectively. Resulting plaques were screened for gain of mRFP-VP26 capsid fluorescence.

PRV1020 and PRV1021 were generated as follows: PRV437 contains a Us9-null mutation and a gM-mCherry fusion, as previously described (41). Shuttle plasmid pSB25 (L. Pomeranz, S. Bratman, and L. Enquist, unpublished data) contains the PRV UL10 gene, encoding glycoprotein M, with a C-terminal EGFP fusion. Plasmid pßactin-PEX3-mRFP-FKBP was a kind gift from C. Hoogenraad (Utrecht U., Netherlands) (37). The 2xFKBP domains from pβactin-PEX3-mRFP-FKBP were amplified by PCR and ligated into pSB25 using SphI/AleI restriction sites, replacing EGFP. The resulting gM-FKBP fusion junctions are as follows (C-terminal gM sequence is in bold type, peptide linkers are italicized, 2xFKBP sequence is in plain type, and stop codon indicated with an asterisk):

**Table 1.**
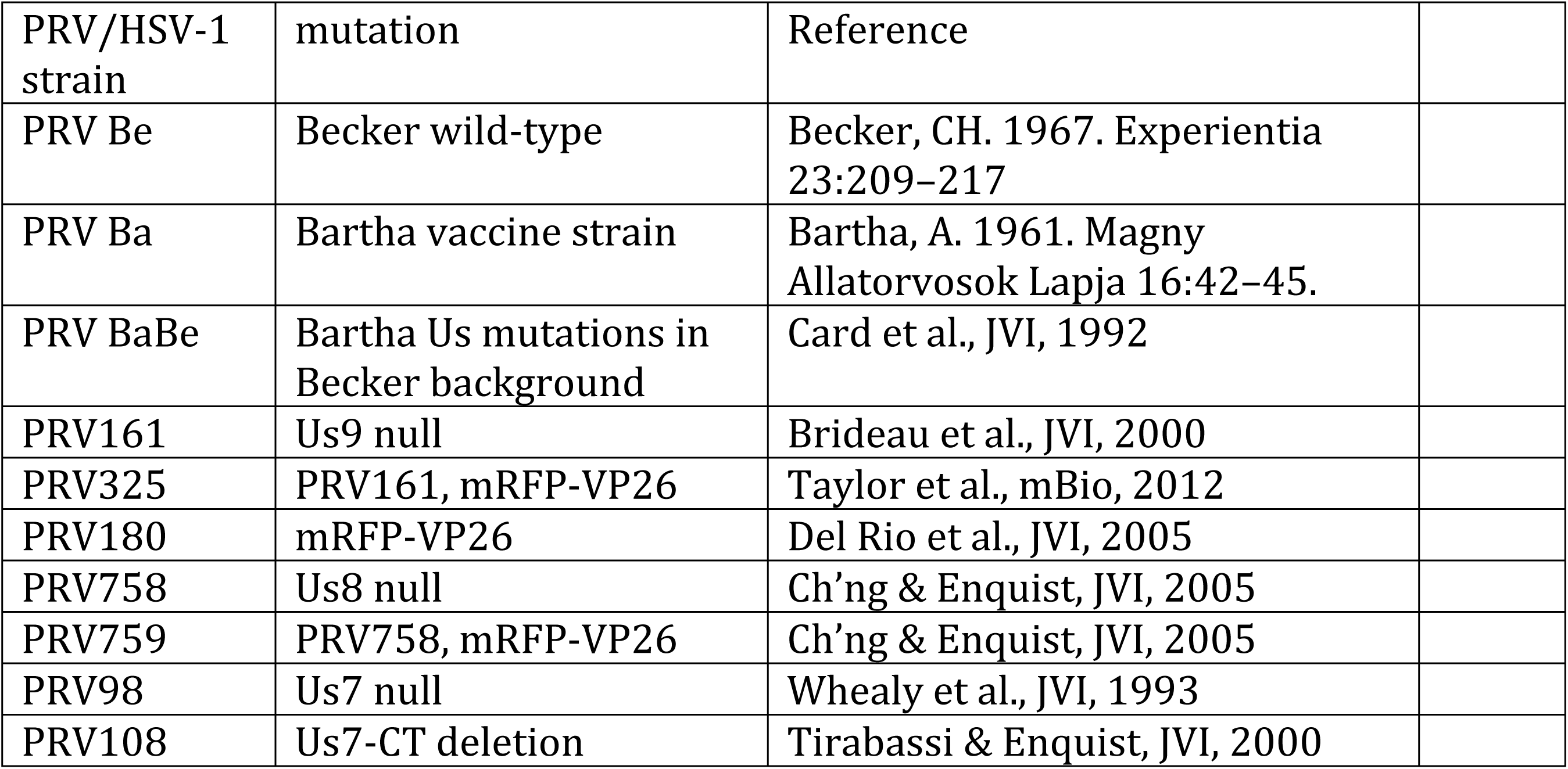
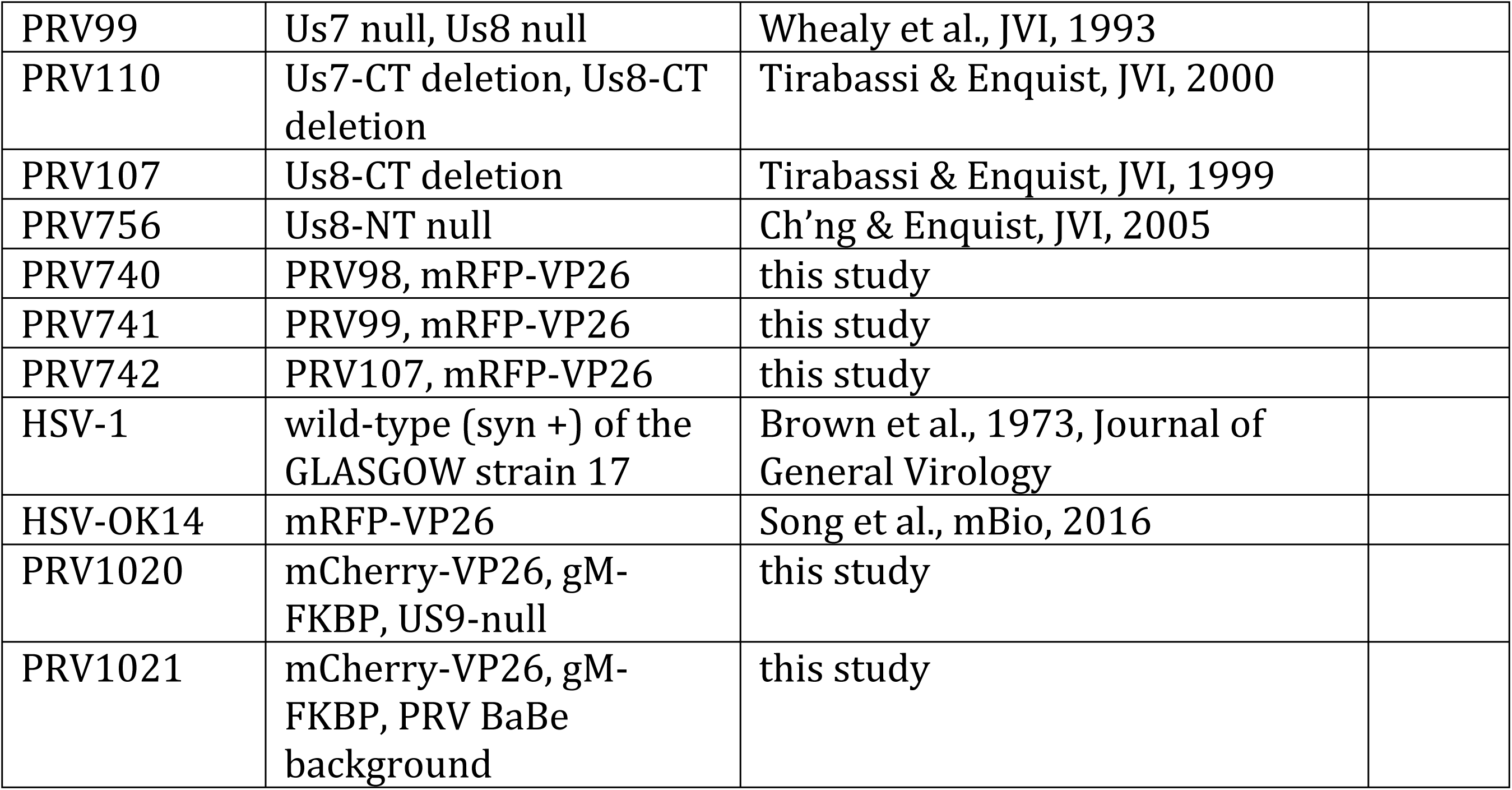
Recombinant PRV and HSV-1 strains used in this study.

…**EVVYENLGFE***MVSKGEELFTD*RGVQVETISP…FDVELLKLET*SYLYK*\*

The gM-FKBP shuttle plasmid was co-transfected with PRV437 nucleocapsid DNA, and plaques were screened for the loss of gM-mCherry fluorescence, producing a cloning intermediate, PRV1024. A shuttle plasmid encoding mCherry-VP26 capsid fusion (42) was co-transfected with PRV1024 nucleocapsid DNA, and plaques were screened for gain of mCherry capsid fluorescence. The resulting PRV1020 expresses an mCherry-VP26 capsid tag, gM-FKBP fusion protein, and contains a US9-null mutation. PRV BaBe consists of wild-type PRV Becker with the Bartha Us region containing a deletion that disrupts genes Us7, Us8, Us9, and Us2. Using the same procedure described above we also introduced the mCherry-VP26 capsid tag and gM-FKBP fusion protein into PRV BaBe, producing PRV1021.

### Adenovirus Vectors

The adenovirus vectors expressing GFP-Us9 and Us8-mCherry were previously described (17, 43). Adenovirus vectors expressing herpesvirus Us7 (gI) or Us8 (gE) were constructed by PCR amplification of the coding sequence from PRV strain Becker or HSV strain 17. PRV Us9 wild-type and mutant genes were synthesized (Genscript, Piscataway, NJ). The cellular gene Kif1A was PCR amplified from rat cDNA. The plasmid pβactin-KIF1A(1–489)-GFP-FRB was a kind gift from C. Hoogenraad (Utrecht U., Netherlands) (37). All genes of interest were cloned into pAd/CMV/V5-DEST by Gateway recombination (Invitrogen). The completed vectors were confirmed by DNA sequencing. All adenovirus vectors were propagated on complementing 293FT cells, cell-associated virus was harvested in serum-free DMEM, and the transduction efficiency of the resulting stocks was estimated by visualizing fluorescent protein expression in rat2 cells. Cell lines were generally transduced for 24 hours.

### Culture of Primary Neuronal Cells

Neurons of embryonic rat superior cervical ganglia (SCG) were isolated and cultured as previously described (13). Briefly, SCGs were dissected from embryonic day 17 (E17) Sprague-Dawley rat embryos (Hilltop Labs Inc., Pittsburgh, PA) and then plated and maintained in neuronal medium consisting of Neurobasal medium (Invitrogen) supplemented with 1% penicillin-streptomycin with 2 mM glutamine (Invitrogen), 2% B27 (Invitrogen), and 100 ng/ml NGF (NB+ medium). Plates were either 35mm plastic dishes (Fisher Scientific, Allentown, PA) for spread experiments, or for live cell analysis MatTek glass-bottom (Ashland, MA) or plastic-bottom (IBIDI, Madison, WI) dishes, all pre-coated before cell plating with 500 μg/ml of poly-DL-ornithine (Sigma-Aldrich, St. Louis, MO) for 24 h at 37 °C and 10 μg/ml of natural murine laminin (Invitrogen) for at least 12 h at 37 °C. Plastic dishes were further prepared for chambered SCG cultures, to separate axon extensions from cell bodies, by mounting modified Campenot chambers (Figure 1 A). A series of parallel grooves were etched across the surface and covered with 1% methylcellulose in DMEM. A CAMP320 three-chambered Teflon ring (Tyler Research; Edmonton, Alberta, Canada) was then coated with vacuum grease on one side and placed on top of the tissue culture surface, oriented such that the grooves extended across all three compartments. To kill non-neuronal dividing cells, 48 h after SCG cell plating, 1 mM cytosine-D-arabinofuranoside (Sigma-Aldrich) was added for 2 days. All SCG cultures were allowed to differentiate for at least 14 days prior to infection.

All animal work was performed in accordance with the Princeton Institutional Animal Care and Use Committee (protocol #1947)

### Anterograde Spread Assay and Herpesvirus Infections

Chambered SCG cultures were transduced in the soma compartment with adenovirus vectors 2-3 days prior to herpesvirus infections. Transduction efficiency of vectors was checked by fluorescence microscopy to be >95 % before 10^5^ indicator cells were plated in the neurite compartment and 1 % methocel in NB+ medium was added to the middle compartment to further prevent medium exchange between chambers. Chambered SCG cultures were inoculated in the S compartment with either PRV or HSV with 10^6^ pfu (∼MOI of 50) to achieve synchronized infections. To determine titers, soma and neurite chamber media were harvested 24 hours post-infection (hpi)., if not stated otherwise. Live microscopy of entire chambers was initiated between 6-8 hpi and lasted up to 72 h. Dissociated SCG cultures were also infected with 10^6^ pfu (∼MOI of 50) in a small volume of NB+ medium.

PRV titers were determined by plaque assay on PK15 cells overlaid with 1% methocel in DMEM containing 2 % FBS, whereas VERO cells were used for HSV strains. Brefeldin A (BrefA) was purchased from Sigma-Aldrich (B7651) and added to the cells at a concentration of 5 μg/ml.

### Biochemical Methods, Western Blotting, and Antibodies

Detergent resistant membrane (DRM) fractions were isolated by Optiprep sucrose gradient centrifugation as previously described (34). Briefly, ∼10^6^ SCG neuronal cells were either mock infected or infected at MOI=10 with PRV Becker or PRV 161. At 12 hpi, cells were collected in a 50 ml conical tube, and washed twice with cold RPMI medium by brief centrifugation at 3,000 rpm. Cells were lysed with 1 ml of lysis buffer consisting of 1% TX-100 in TNE buffer (25 mM Tris HCl [pH 6.8], 150 mM NaCl, 5 mM EDTA), protease inhibitor cocktail (Roche Diagnostics GmbH, Mannheim, Germany), and 5 mM iodoacetamide. The lysate was homogenized by being passed 15 times through an 18-gauge needle, and then allowed to rock for 30 minutes at 4°C. At the end of the rocking period, the sample was again homogenized briefly, and then mixed with 2 ml of ice-cold 60% Optiprep™ density gradient medium (Sigma-Aldrich). The entire 3 ml mixture was placed at the bottom of a Beckman SW41 ultracentrifuge tube (Beckman, Munich, Germany) and subsequently overlaid with 5 ml of ice-cold 30% Optiprep in TNE and 4 ml of ice-cold 5% Optiprep in TNE. Samples were centrifuged at 34,200 rpm (200,000×g) at 4°C for 20 hours. Twelve fractions were collected from the top to the bottom of the tube (1 ml each), and processed for immunoblotting. For direct cell lysate analysis or immunoprecipitations, cells were lysed after two washes twice with PBS (HyClone) and one wash with radioimmunoprecipitation assay (RIPA) buffer (10 mM TrisHCl, 150 mM NaCl, 1 mM EDTA, and 1 mM EGTA, pH7.4). Samples were then taken up in RIPA, kept on ice for 15 min, centrifuged at 13,200 rpm at 4°C, transferred into new tubes and either immediately processed for immunoblotting or incubated for coimmunoprecipitations with monoclonal anti-HA antibodies linked to Sepharose A beads at 4°C for 1.5 h and washed, and the bound (pellet) fractions were analyzed by immunoblotting. Samples were heated in Laemmli buffer at 100°C for 5min before proteins were separated on gradient polyacrylamide gels (4-12%, NuPage, Invitrogen), which were run at 175V for 50min. Proteins were transferred onto a PVDF membrane (0.45 μm pore size, Immobilon-P, Millipore) using semidry transfer (Biorad). After transfer, membranes were incubated in 5% non-fat dry milk in PBST (PBS + 0.1% Tween-20) solution for 1 h at room temperature. Immunoblots were performed using primary and secondary antibodies in 5% milk PBST solution. Membranes were incubated with chemiluminescent substrates (Supersignal West Pico or Dura, Thermo scientific). Protein bands were visualized by exposure on HyBlot CL (Denville scientific) blue X-ray films.

Antibodies used in this study included a monoclonal anti-GM130, anti-TGN38 (both from BD Transduction Laboratories), anti-Kif5 (MAP1614, Chemicon), anti-β-actin (Sigma-Aldrich), anti-transferrin receptor (TfR, Zymed, San Francisco, CA), pooled anti-PRV-gE (kind gift from T. Ben-Porat) and phospho-specific Us9 antibody 2D5E6 (20). Rabbit antibodies included anti-Kif1a (HPA004831 & HPA005442, Sigma-Aldrich), anti-Us9 (22), anti-PRV-gI and anti-PRV-gE (generous gifts from K. Bienkowska-Szewczyk) and goat anti-PRV-gB (284) and anti-PRV-gI proteins were described elsewhere (44).

### Microscopy and Analysis of Capsid Motility

For fixed cell microscopy, dissociated SCG neurons were grown on MatTek dishes (Ashland, MA) for 2 weeks prior to imaging. Upon transduction with adenovirus constructs, cells were fixed in a 4% paraformaldehyde-PBS solution for 10 minutes at room temperature followed by two PBS washes. To label actin filaments for fixed cell analysis, 1 μM SiR-actin (CY-SC001, Cytoskeleton, Denver, CO) was added to SCG cultures one hour prior to fixation. Prior to antibody staining, samples were permeabilized with 0.05% Triton-X in PBS for 10 min at RT followed a PBS wash and blocked with 3% BSA-PBS solution for 1hr at RT followed by five PBS washes. Samples were stained with anti-GM130 antibody raised in rabbit (Ab52649) at 1:750 dilution followed by 2X PBS washes, then incubated with Alexa-546 secondary antibody for 1hr RT followed by a final wash in PBS. For confocal microscopy, images were acquired on a Nikon A1 confocal microscope with a Nikon 60x oil objective (NA = 1.4). For super-resolution microscopy, 3D structured illumination microscopy (3D-SIM) was performed using a Nikon N-SIM. All data for 3D-SIM were captured used a Nikon CFI Apo TIRF (100× 1.49 NA oil objective). 3D-SIM image stacks were sectioned using a 250 nm z-step size. Raw image data were reconstructed and analyzed using Nikon Elements software.

Live anterograde spread in chambered SCG cultures was imaged using a Nikon Ti-Eclipse inverted epifluorescence microscope (Nikon Instruments, Melville, NY) equipped with a separate fast-switching excitation and emission filter wheels (Prior Scientific, Rockland, MA) and a heated cell culture chamber (Ibidi) to ensure biologically relevant environmental conditions during image acquisition. Images were acquired every two hours with a CoolSnap ES2 charge-coupled device (CCD) camera (Photometrics, Tucson, AZ) and a Nikon Plan Fluor PhlDlWD 4× (N.A. 0.13) objective (Nikon Instruments) in large image mode covering the entire chamber. For microscopy of axonal transport in neuronal cells, we used high resolution dual-color imaging on an inverted Nikon Ti-Eclipse TIRF microscope equipped with an Agilent laser source (Agilent) producing approximately 65 mW of 488-nm and 561-nm light at the fiber exit, an Nikon APO TIRF 100× (N.A. 1.49) objective, and an iXon Ultra EM-CCD camera (Andor Technology, South Windsor, CT), run through Nikon NIS AR software (Princeton University Molecular Biology Confocal Microscopy Facility). Lasers at 405, 488, and 561 nm were used at no more than 30% of their maximal output and allowed trigger acquisition at a frame rate of 18.7 fps (two colors) or 15.7 fps (three colors) with 25 nm spatial resolution.

Particle tracking and motility analysis was performed as described earlier for adenovirus (45), but modified for alpha-herpesviruses (13). Briefly, a custom-tracking algorithm was used to extract the position of virus particles as a function of time. All fluorescent particles were fit to 2-D Gaussian profiles and treated as coming from a diffraction-limited point source. Positions of each particle were obtained from the Gaussian fits. Trajectories were reconstructed by matching particle locations based on their displacements between frames. To limit contributions of rare but long non-motile particle tracks as well as to reduce the relative contribution of very long pauses, parts of tracks with asymmetry parameter below 1.0 (±25 point window) were eliminated from analysis. Trajectories were then automatically parsed into segments of constant velocity (see also Figure 4). Previously published analyses focused on motile events longer than 400 - 500 nm. To mimic this, velocities were analyzed for long segments above the standard motility cut-off only (2σ value of the zero-centered peak in segment length distribution). The cut-off was usually in the range of two to four capsid diameters (300 - 600 nm). All segments above the cut-off were used to assemble segment length histograms with decay curve fits.

### Ethics Statement”

All animal work was conducted in accordance with the Princeton Institutional Animal Care and Use Committee (protocol # 1947). All personnel adhered to applicable federal, state, local, and institutional laws and policies governing ethical animal research. This includes the Animal Welfare Act (AWA), the Public Health Service Policy on Humane Care and Use of Laboratory Animals, the Principles for the Utilization and Care of Vertebrate Animals Used in Testing, Research and Training, and the Health Research Extension Act of 1985.

## Acknowledgements

We are grateful to G. Laevsky at the confocal imaging facility for his help with the confocal, TIRF, and 3D-SIM microscopes (Department of Molecular Biology, Princeton University), as well as all other former and current members of the Enquist laboratory for helpful discussions.

---

**Supplemental Figure 1.**
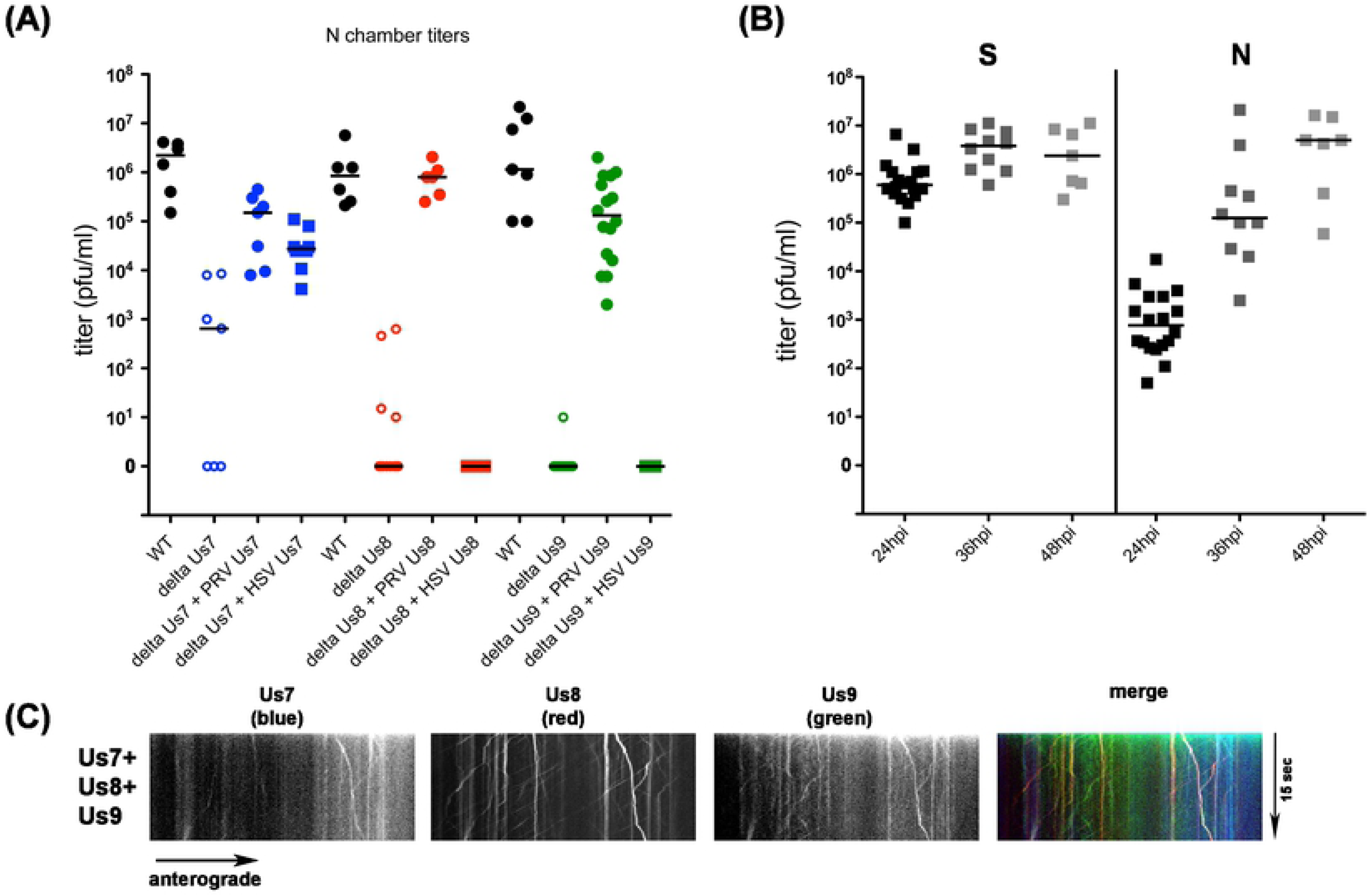
(A) Titers of PRV single Us7, Us8, or Us9 gene deletion mutants 24 hpi and rescue by adenovirus transduction with according PRV or HSV-1 gene produce. Effects on anterograde spread is indicated by N compartment titers. Each data point represents one chamber; horizontal bars indicate median values for each condition. Only HSV-1 Us7 is able to rescue PRV ΔUs7 virus, whereas HSV-1 Us8 and Us9 fail to rescue PRV ΔUs8 or ΔUs9, respectively. (B) Titers of HSV-1 wild-type infected anterograde spread assays at 24h, 36h, and 48h post-infection. Titers left and right of the solid vertical line represent S and N compartment titers, which indicate replication and anterograde spread, respectively. Each data point represents one chamber; horizontal bars indicate median values for each condition. Compared to PRV (Figure 1), HSV-1 has an up to 24h delayed anterograde spread phenotype. (C) TIRF microscopy of live SCG axons grown in compartmentalized cultures. Cells were transduced with HSV-1 proteins Us7, Us8, and Us9 and imaged at ∼12frames/s in three-color mode. Axonal co-transport of Us7-9 was observed without HSV-1 infection.

